# Maternal *schistosomiasis* impairs offspring IL-4 production and B cell expansion

**DOI:** 10.1101/2020.08.13.250274

**Authors:** Diana Cortes-Selva, Lisa Gibbs, Andrew Ready, H. Atakan Ekiz, Bartek Rajwa, Keke C. Fairfax

**Affiliations:** Department of Pathology, Division of Microbiology and Immunology, University of Utah, Salt Lake City UT, 84112; Bindley Bioscience Center, Purdue University, West Lafayette IN, 47907; Department of Comparative Pathobiology, College of Veterinary Medicine, Purdue University, West Lafayette IN, 47907

**Keywords:** Maternal infection, *Schistosoma mansoni*, memory T cells, memory B cells, Interleukin-4, follicular dendritic cells

## Abstract

Maternal helminth infections are a global public health concern and correlate with altered infant immune responses to some childhood immunizations, but a mechanistic understanding of how maternal helminth infection alters the cellular immune responses of offspring is lacking. Here we establish a model of maternal *Schistosoma mansoni* infection in dual IL-4 reporter mice. We find that offspring born to mothers infected with *S. mansoni* have impaired production of IL-4 during homoeostasis, and following immunization with a Tetanus-Diphtheria vaccine. We identified that iNKT cells are the dominant source of IL-4 during early life homeostasis, and that diminished IL-4 production was associated with both reduced B cell and follicular dendritic cell responses. These defects were maintained long-term, affecting memory B and T cell responses. Single-cell RNASeq analysis of immunized offspring identified egg antigen-dependent reductions in B-cell cell cycle and proliferation-related genes. These data reveal that maternal infection leads to long-lasting defects in the cellular responses to heterologous antigens and provide vital insight into the influence of maternal infection on offspring immunity.

## Introduction

Schistosomiasis is an infectious disease caused by trematode parasites of the genus Schistosoma, with *S. mansoni, S. haematobium* and *S. japonicum* causing the most human morbidity [1]. This disease affects an estimated 779 million people who are at risk, and 207 million people infected annually [2], with the highest prevalence in adolescents and young adults (10 to 24 years of age) [3]. This leads to a unique impact on women of reproductive age, with 40 million women infected annually. Human studies on schistosomiasis and pregnancy have previously established a link between maternal infection and low birth weight, as well as premature birth and intrauterine growth restriction [4-6]. Moreover, examination of the effects of maternal schistosomiasis on the response of offspring to heterologous antigens has demonstrated impaired responses in childhood-vaccine induced immunity, raising the concern that a mother’s parasitic status during pregnancy might render early childhood immunization ineffective for years and even decades post-immunization [7]. Decreased responses to bacillus Calmette-Guérin (BCG) have been shown in children sensitized *in utero* to *Schistosoma haematobium* [8]. Furthermore, children born to mothers infected with *S. mansoni* presented lower anti-measles antibody levels [7] at 2 years of age. Altogether, these data suggest that populations where schistosomiasis is endemic, are vulnerable to vaccine failure, and as a consequence, susceptible to deaths due to preventable infectious diseases. Endemic regions may require an adjusted immunization regimen in order to ensure optimal protection against diseases, but mechanistic studies are needed to examine this.

Similar to the human data, mouse studies have shown reduced protection from BCG vaccine against *Mycobacterium tuberculosis* challenge in mice infected with *S. mansoni [9]*, while infection during gestation has shown increased susceptibility of offspring to subsequent *S. mansoni* infection [10]. Many authors have described a state of hyporesponsiveness to homologous antigens that has been attributed, in part, to immune factors acquired from the mother and/or previous antigen contact [11]. Previous sensitization of murine offspring to antigens from mothers infected with other parasitic infections such as *Wuchereria bancrofti* and/or *Plasmodium falciparum* have shown a Th-2 biased response (higher production of *il-4* mRNA and IgE antibody) to diverse antigens when compared to mice from uninfected mothers [12]. However, the effects of schistosomiasis during pregnancy on the immune response in offspring during homeostasis, and when challenged with heterologous antigens remain poorly defined [13]. *S. mansoni* infection elicits host responses that are similar in humans and mice, so the murine model of schistosomiasis is useful to determine the mechanisms by which prenatal infections induce diminished postnatal immune responses.

In this study we aimed to define the effects of perinatal *S. mansoni* infection on offspring homeostatic immunity during early life, and determine whether maternal infection causes a deleterious or beneficial effect in response to immunization with heterologous antigens. We have established a novel experimental murine model of maternal schistosomiasis in IL-4 dual reporter mice. Our data from this model demonstrate that maternal schistosomiasis reduces steady-state iNKT and CD4-T cell production of IL-4. This reduction correlates with reduced follicular dendritic cells (FDCs) and circulating plasma and memory B cells at steady state Additionally, the cellular responses to immunization with commercially licensed tetanus/diphtheria vaccine are diminished following primary immunization, leading to a defect in memory TFH precursors. Single cell RNASeq analysis revealed reductions in multiple cell cycle/proliferation genes, as well as the critical B cell transcription factor EBF-1. Thus, *in utero* exposure to *S. mansoni* egg antigens induces long-lived modulation of the development of humoral and cellular responses via suppression of IL-4 production, and the B cell-stromal cell axis.

## Results

### Maternal *Schistosoma mansoni* infection results in *in utero* egg antigen sensitization and a reduction in the B cell-stromal cell axis

We infected 4get homozygous (IL-4 reporter mice on a BALB/c background [14]) female mice with a low dose of *S. mansoni* (35 cercariae) as described in Materials and Methods. At five to six-weeks post infection (the timepoint where egg production begins) they were paired with a naïve KN2 homozygous (IL-4 deficient [15]) male. At 6 weeks post infection females were tested for antibodies specific to Schistosoma egg antigen (SEA) to confirm patent infection. Age-matched mixed-gender pups from infected and control uninfected mothers were weaned at 28 days to maximize survival and lactation period. At 28-35 days old, pups were sacrificed for steady-state studies. Egg antigen (SEA) specific antibodies in pups from infected and uninfected mothers were tested at 35, 90 and 180 days of age (Figure 1 A). As expected, pups from mock-infected (controls) showed no antigen-specific response to SEA at any of the timepoints. In contrast, pups from infected mothers exhibited an anti-SEA IgG1 antibody response at 35, 90 and 180 days of age. In some cases, at 180 days of age, some of the pups exhibited no detectable anti-SEA IgG1 (Figure 1 A), indicating heterogeneity in the humoral response to egg antigens. As expected, we found that at 28-35 days of age, offspring anti-SEA IgG1 titers correlate with maternal titers (Figure 1B, p<0.0001 R^2^=0.902). Next, we wondered if the antigen-specific antibody was maternally derived or self-derived. For this, we mated 4get/KN2 (infected or uninfected) females and KN2 homozygous males to obtain KN2 homozygous or 4get/KN2 littermate offspring. Since KN2 homozygous mice are IL-4 deficient, they should have a diminished ability to switch to IgG1 in response to SEA [16]. We observed that at 35, 60 and 90 days of age KN2 and 4get/KN2 homozygous had detectable anti-SEA IgG1 antibodies, and that titers were similar between KN2 and 4get/KN2 animals (Figure 1 C).

**Figure 1.**
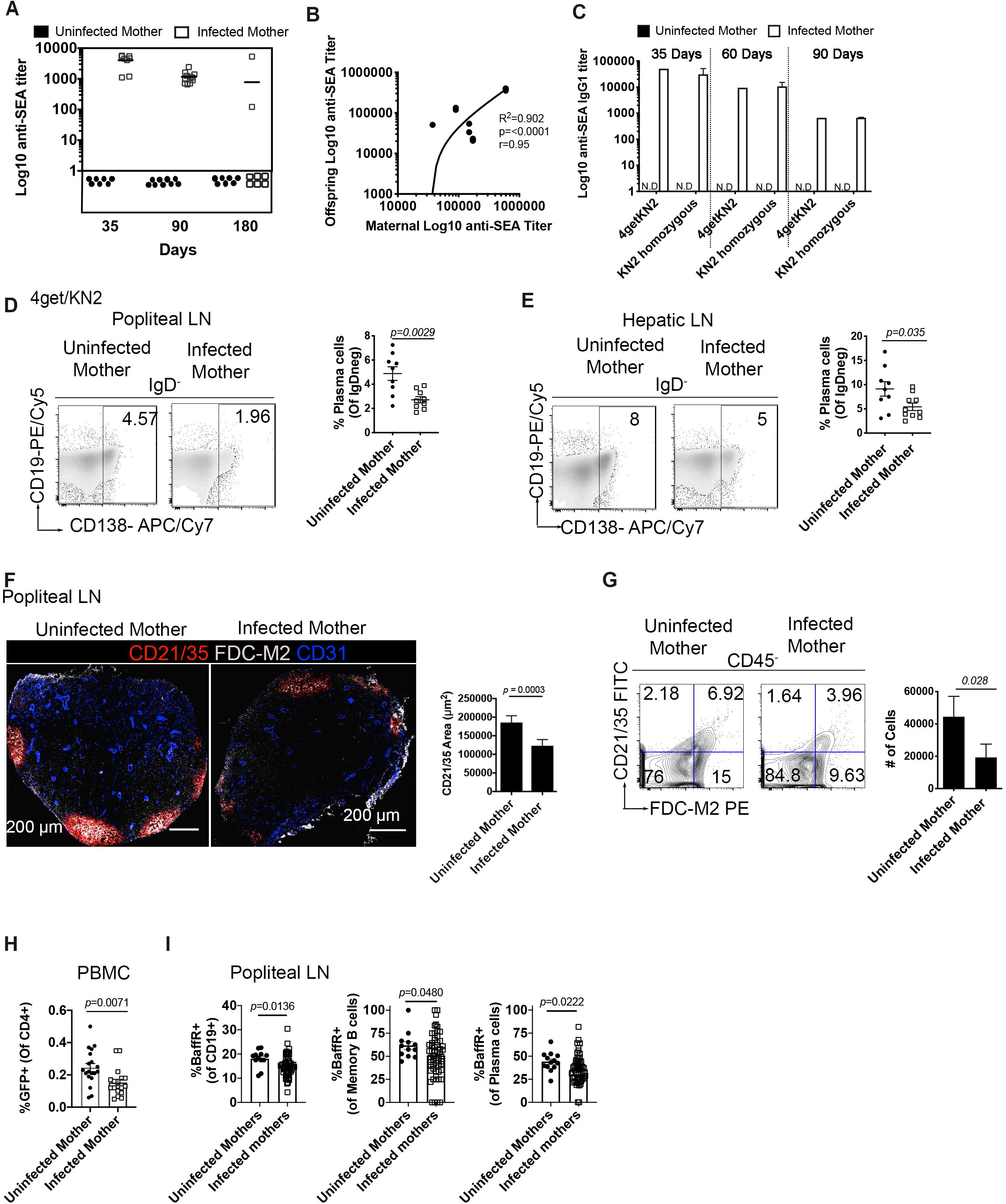
Maternal *S. mansoni* infection leads to a humoral anti-SEA response in offspring and reduction in follicular dendritic cells. (A) anti-*Schistosoma mansoni* egg antigen (SEA) specific IgG1 antibody titers in serum of naïve 4get/KN2 mice born to infected or uninfected mothers. Each point represents an individual mouse. (B) Analysis of the correlation between maternal and offspring anti-SEA specific IgG1. (C) Anti-SEA IgG1 titers in naïve KN2 homozygous and 4get/KN2 mice born to infected and uninfected mothers at 35, 60 and 90 days of age. (D-E) Flow cytometry plots of plasma cells (IgD^-^CD19^+/-^CD138^+^) in popliteal and hepatic lymph nodes from naïve 4get/KN2 pups born to infected and uninfected mothers. (F) Tile confocal imaging of naïve popliteal lymph nodes of 4get/KN2 mice at 28-35 days of age, scale bar: 200 μm. Sections were stained for CD21/35 (red) FDC-M2 (gray) and CD31 (blue) with the respective CD21/35 area quantification. (G) flow cytometry analysis of FDCs (CD21/35^+^FDC-M2^+^) gated from CD45^-^ with total number of FDC in popliteal lymph node from pups 28-35 days of age. (H) Frequency of CD4^+^ GFP^+^ in the blood of naïve 4get/KN2 mice. Flow plots were concatenated samples, with n >3 mice per group and representative of six independent experiments. Confocal microscopy data is representative of n >3 mice per group from two biologically independent experiments. Statistical significance was calculated by unpaired Student’s t□test. Error bar denotes mean ± SEM. Correlation analysis was calculated by Pearson correlation coefficient.

Exposure *in utero* to *S. mansoni* egg antigens correlated to a significantly diminished bulk plasma cell population at steady-state in both popliteal (Figure 1D) and hepatic lymph nodes (Figure 1E), with a 42% reduction of plasma cells in popliteal lymph nodes and a 60% reduction of total plasma cells in hepatic lymph nodes. Follicular dendritic cells trap immunocomplexes via Fc and Complement receptors [17]. Previous studies have established that FDCs serve as sites of B cell antigen capture [18], and that persisting antigens trapped by FDC induce somatic hypermutation [19]. Since we found that pups born to infected mothers have reduced plasma cells in lymph nodes, we sought to determine whether this is accompanied by modifications in the stromal cell population of follicular dendritic cells (defined as CD21/35, complement receptor 2 and 1, respectively and FDC-M2 positive) via tile confocal microscopy. Indeed, in pups from infected mothers, a marked decrease of intensity in the markers for FDCs is observed (Figure 1 F). In addition to reduced intensity of CD21/35 and FDC-M2 in naïve peripheral lymph nodes, we also observed reduced area of CD21/35 FDCM2 double-positive cells. To confirm the confocal tile scan results, we analyzed lymph nodes by flow cytometry and confirmed that offspring born to *S. mansoni* infected mothers have reduced frequency and total cell number of FDC in peripheral lymph nodes (Figure 1 G). We then analyzed the lymphocyte compartment to determine if B or CD4 T cell development is altered by maternal *S. mansoni* infection. We found no difference in the absolute numbers of CD19 cells in peripheral lymph nodes (Data not shown). Our recent work demonstrated a role for IL-4 in the homeostatic maintenance of FDCs [20], so we analyzed IL-4 competent Th2 CD4-T cells (GFP^+^) and found a reduction in Th2 cells in the blood (PBMCs Figure 1 H). The Baff-BaffR signaling pathway is known to be critical for long-term B cell survival, so we measured BaffR expression via FACS and found reduced BaffR in bulk CD19+, plasma, and memory B cells (Figure 1I). These data suggest that egg antigen sensitization alters two critical pathways that provide survival signals to B cells, BaffR and immune complex presentation.

### Invariant NK T cells are the source of IL-4 in the peripheral lymph node at steady state

Since we observed reduced Th2 (GFP^+^) CD4 T-cells at steady-state in the blood of pups from infected mothers in comparison to control pups, we analyzed a diverse spectrum of cells, including gamma delta T cells, alpha beta T cells, NK, NK T cells and iNKT cells to look for the steady-state secretion of IL-4 (marked by HuCD2). We observed very little secretion of IL-4 in any cell populations except for invariant NKT-cells (Figure 2 A). Moreover, there was a marked decrease in both transcription and secretion of IL-4 in the lymph nodes of offspring from infected mothers, with a 5-fold reduction in IL-4 secreting iNKT cells from pups that were antenatally exposed to SEA antigens (Figure 2 B). These data suggest that steady-state IL-4 production is attenuated by antenatal *S. mansoni* antigen exposure, and establishes that iNKT cells are the homeostatic source of lymph node IL-4 during early life.

**Figure 2.**
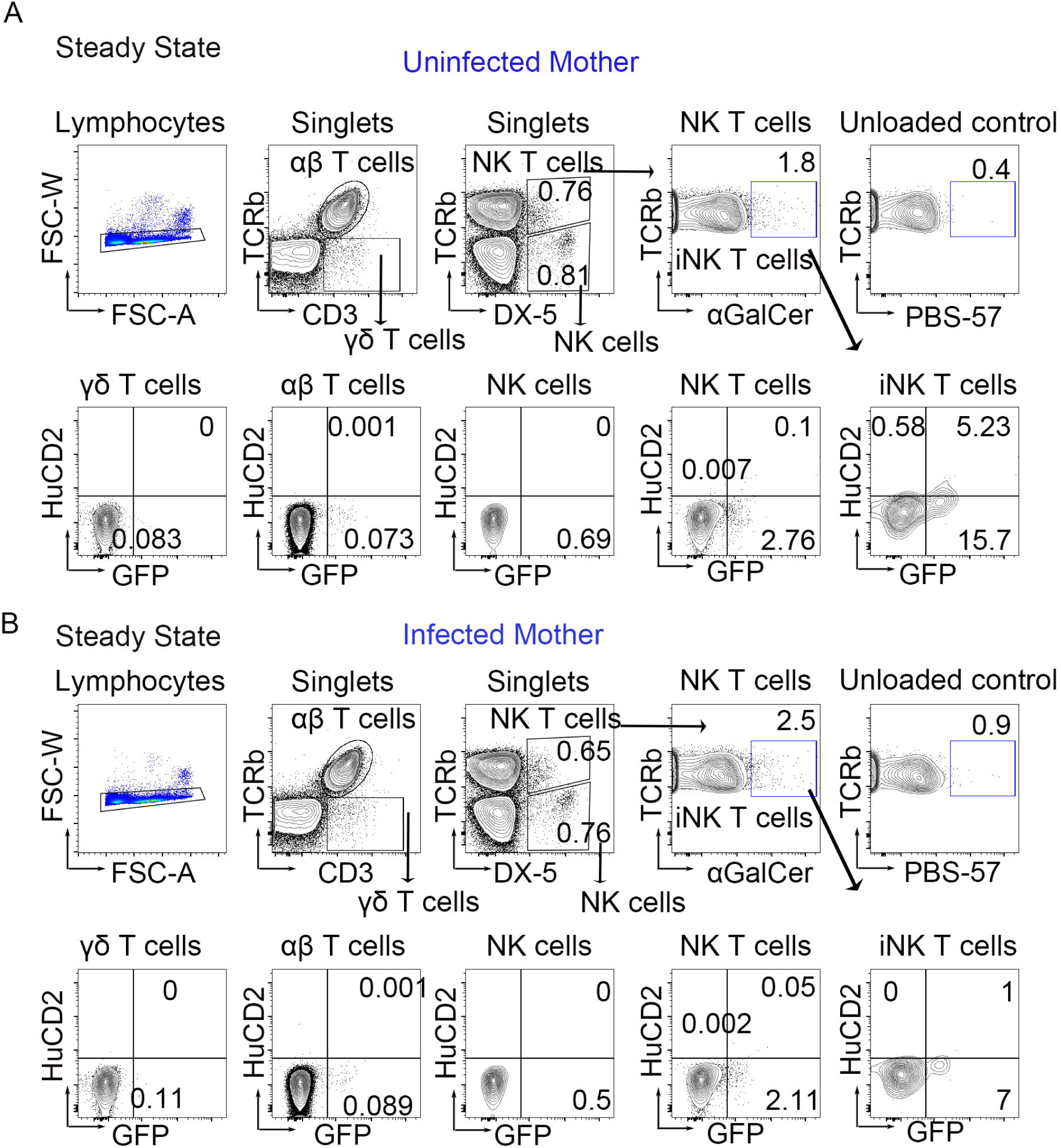
iNKT cells are the main cellular source of IL-4 in peripheral lymph nodes at steady state, and maternal *S. mansoni* infection reduces their secretion of IL-4. Gating strategy in popliteal lymph nodes for invariant NKT cell (αGalCer^+^TCRαβ^+^ DX-5^+^), αβ T cells (TCRβ^+^CD3^+^), γδT cells (TCRβ^-^CD3^+^), NK T cells (TCRβ^+^DX-5^+^), NK cells (TCRβ^-^DX-5^+^) with respective staining control (unloaded tetramer) analyzed by flow cytometry in (A) pups from uninfected mother and (B) pups from infected mother at 28-35 days of age. Flow plots are concatenated from >4 mice per group with two biologically independent experiments.

### Offspring born to infected mothers have reduced Th2 development following primary immunization with tetanus/diphtheria

Having confirmed that maternal infection induces changes in the immune response in offspring at steady state, we wondered whether the immune response to heterologous antigens could be affected by *S. mansoni* maternal infection. We immunized 28-35-day-old 4get/KN2 pups subcutaneously (rear footpad injection) from infected and uninfected mothers with ∼1/10^th^ of the human dose of adjuvanted Tetanus/Diphtheria vaccine [20]. At eight days post-immunization we observed that pups from infected mothers exhibited reduced germinal centers in popliteal lymph nodes as well as reduced IL-4 production in the draining lymph node (Figure 3 A). Moreover, offspring born to infected mothers presented significantly reduced IL-4 production within germinal centers (arrowheads indicate HuCD2^+^ CD4^+^ cells, Figure 3 A). In addition to reduced frequency, immunofluorescence analysis revealed that germinal center area was reduced in the draining lymph node of mice born to infected mothers (Figure 3 A). Subsequently, we analyzed the T follicular cells (Tfh) and observed a significant reduction of bulk Tfh cells at 8 days post-immunization in offspring from infected mothers (Figure 3 B), but no change in individual TFH cell production of IL-21, IFNγ, of IL-4 (Figure 3 C,D). This TFH cell reduction was accompanied by reduced effector and memory Th2 responses (Figure 3 E,F). Additionally, mice from *S. mansoni* infected mothers had significantly reduced numbers of germinal center and IgG1^+^ B cells by flow cytometry (Figure 3 G,H,J)). We also observed a defect in the bulk memory B cell response in the pups from infected mother (Figure 3 I, K). This suggests that maternal infection suppresses the cellular immune response to heterologous antigens such as tetanus/diphtheria immunization.

**Figure 3.**
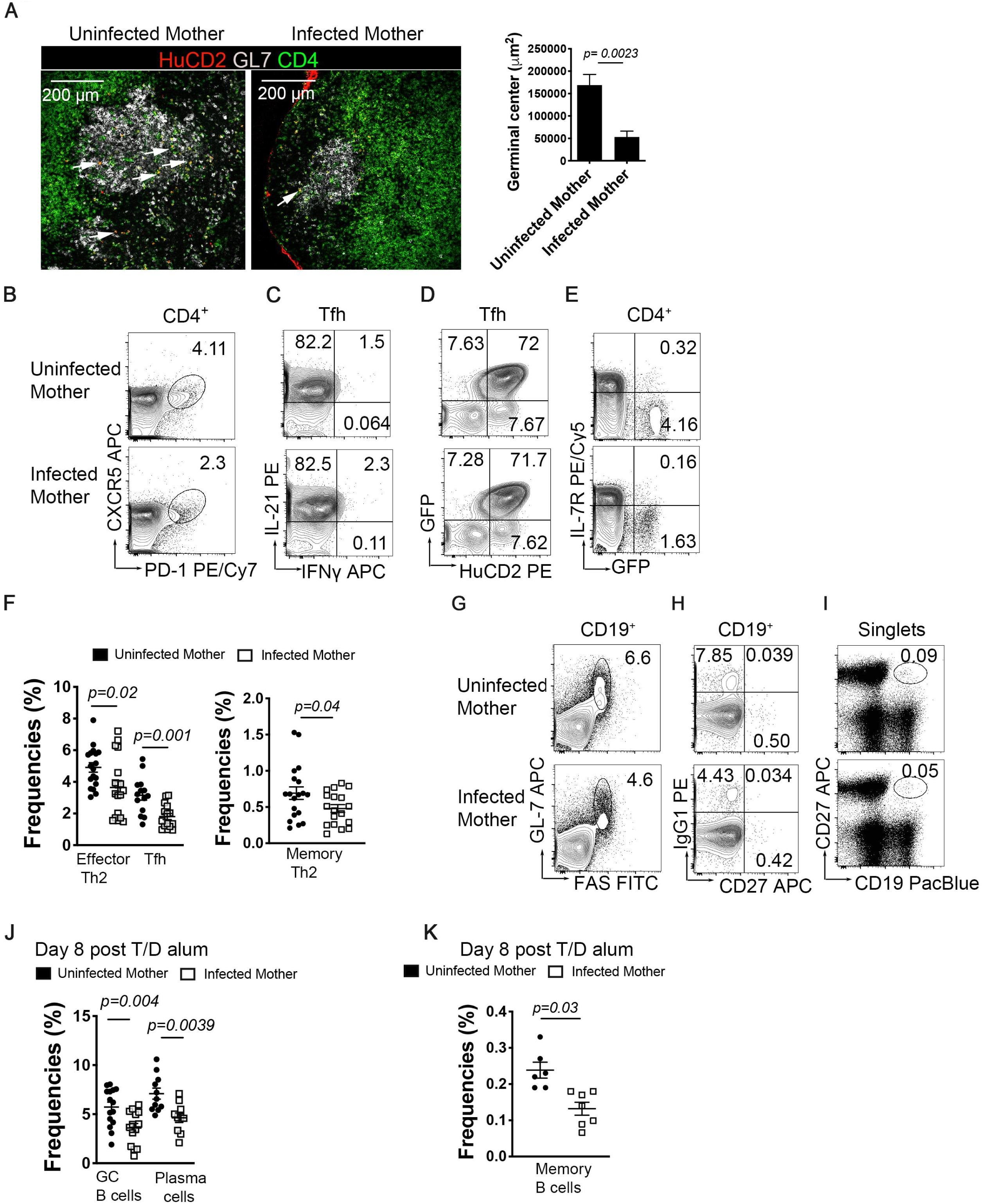
Maternal schistosomiasis leads to reduced TFH cells and IL-4 secretion in response to Tetanus/Diphtheria immunization. 4get/KN2 offspring born to *S. mansoni* infected mothers or uninfected control mothers were immunized in the rear footpad with 1/10^th^ of the human dose of an aluminum adjuvanted Tetanus/Diphtheria vaccine. (A) Tile confocal imaging of 10-micron cryo-sections of popliteal lymph nodes at day-8 post-immunization, stained for HuCD2 (red), GL-7 (gray) and CD4 (green). Scale bar: 200 μm and respective quantitation of germinal center area. Flow cytometry analysis from popliteal lymph nodes of (B) T follicular helper cells (PD-1^+^CXCR5^+^, gated from CD4^+^), (C) intracellular IL-21 and INF-gamma co-production by TFH (D) IL-4 production and secretion by TFH (E) Th2 effector (GFP^+^IL-7R^-^) and Th2 memory (GFP^+^IL-7R^+^). (F) Frequencies of effector Th2, memory Th2 and Tfh cells at 8 days post immunization. (G) Flow cytometry of germinal center B cells (CD19^+^GL-7^+^FAS^+^) (H) Memory IgG1^+^CD27^+^ B cells (I) Bulk memory B cells (CD27^+^CD19^+^) (J) Cell frequencies of GC B cells and plasma cells (K) Frequency of bulk memory B cell in popliteal lymph node. Confocal microscopy images are representative of four independent experiments. Flow plots are concatenated from n >3 mice from at least two biologically independent experiments. Statistical significance was calculated using Student’s t-test.

### *S. mansoni* infection during pregnancy correlates with a weakened immune response in offspring at 14 days post-immunization

In order to determine the effects of antenatal infection on the development of humoral response, we immunized 4get/KN2 pups with a commercial Tetanus/Diphtheria vaccine as above. At 14 days post-primary immunization we observed a defect in Tfh cells (Figure 4 A) that corresponded with a reduced memory and effector Th2 response (Figure 4 B) and decreased plasma cells in pups born to infected mothers (Figure 4 C). Examining the FDC cell compartment, we found that maternal infection leads to decreased antigen-induced FDC expansion and CD21/35 production (quantified by flow cytometry Figure 4 D, E). In addition to reduced frequency, pups from infected mothers exhibited reduced area and mean fluorescence intensity (MFI) of FDCs and germinal centers in draining peripheral lymph nodes (Figure 4 F, G). We then asked if the observed reduction in bulk plasma cells and reduced germinal center area led to reduced anti-tetanus or anti-diphtheria titers. We found no correlation between offspring anti-SEA IgG1 titers and anti-tetanus titers, but did find a correlation (R^2^=0.136, p=.044) between anti-SEA and anti-diphtheria IgG1 titers, suggesting that the diphtheria specific response is more adversely affected by antenatal schistosome exposure than the anti-tetanus response. This is likely tied to the reduction in IL-4 secretion in the germinal center reaction, as our recent work demonstrated that tetanus and diphtheria have a differential dependence on IL-4 for IgG1 class switching [20].

**Figure 4.**
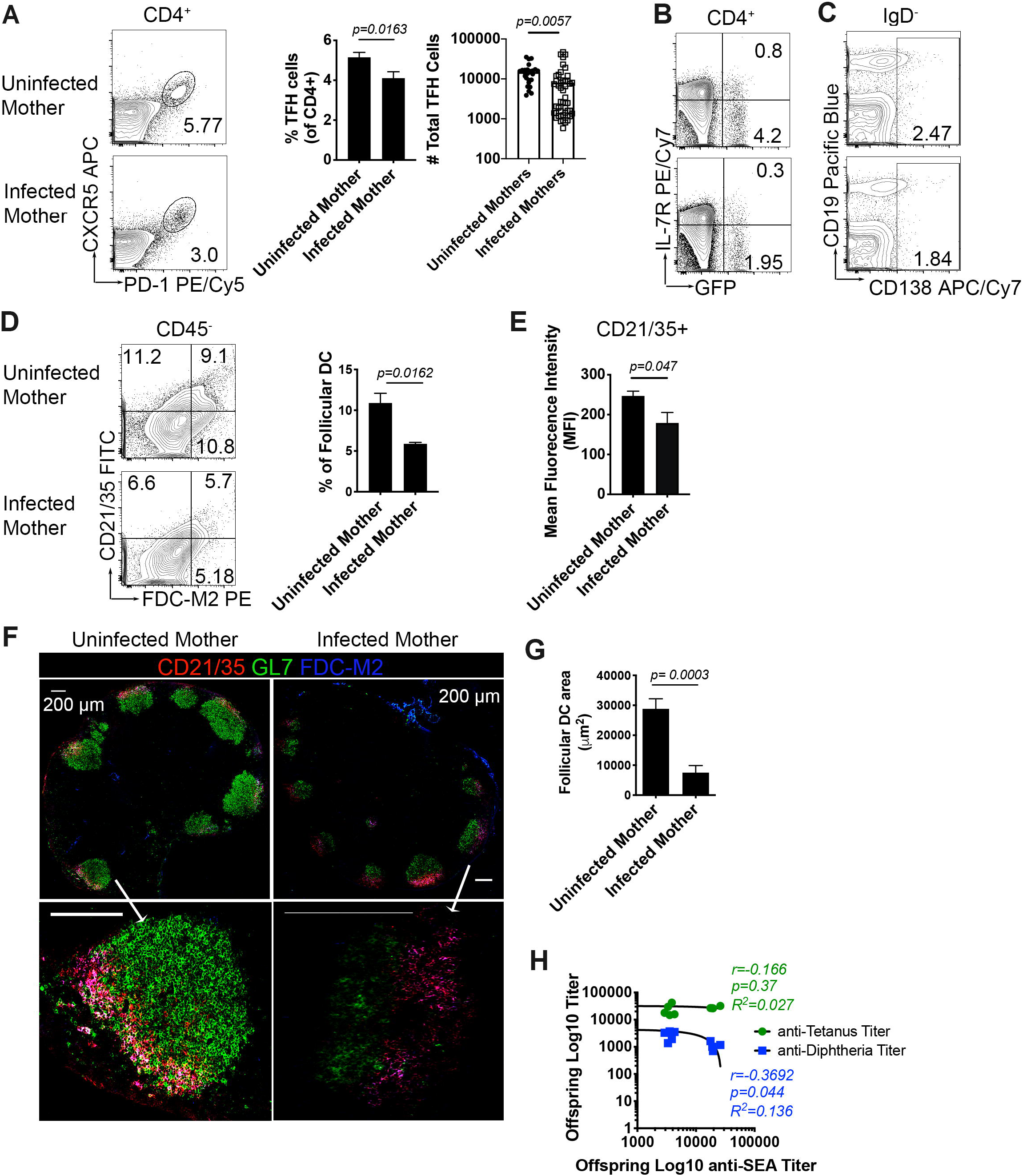
Pups from infected mothers exhibit reduced GC B cells, Tfh, and FDC 14 days post immunization with Tetanus/Diphtheria. (A) Tfh (PD-1^+^CXCR5^+^, gated from CD4^+^) flow plots, frequency and cell number from draining PLN. (B) Th2 effector (GFP^+^IL-7R^-^) and Th2 memory (GFP^+^IL-7R^+^) from reactive lymph node (C) Plasma cells (IgD^-^CD19^+/-^CD138^+^) (D) FDC (CD21/35^+^FDC-M2^+^CD45^-^) frequencies (E) Median fluorescence intensity of CD21/35 on follicular dendritic cells. (F) Cryosections of reactive popliteal lymph node stained with CD21/35 (red), GL-7 (green) and FDC-M2 (blue). Scale bar: 200um. (G) Bar graph of the area of follicular dendritic cells the B cell follicles of draining lymph nodes. (H) Correlation analysis of offspring anti SEA titers to titers of anti-diphtheria and anti-tetanus 14 days post-immunization. Flow plots and confocal microscopy data are representative of three independent experiments with n>3 mice per group. Statistical analysis was calculated with Student’s t-test with Welch’s correction. Correlation was expressed as Pearson correlation coefficient. Error bar denotes mean ± SEM.

### Schistosomiasis during pregnancy alters the memory response to heterologous antigens in the offspring

Immunological memory is key to a protective immune response. Our previous work has demonstrated that subcutaneous immunization induces a persistent germinal center that continues to generate antigen-specific B cells at a low level, and that these memory B cells are critical for an accelerated secondary immune response [16]. Examination of the response to tetanus/diphtheria during maternal schistosomiasis revealed that at 60-90 days post immunization, mice born to infected mothers exhibited a defect in germinal center persistence (Figure 5 A) accompanied by reduced Tfh cells and the absolute number of memory TFH precursors (CXCR5^+^PD1^-^) (Figure 5 B). The Th2 response (IL-4 transcription) was also significantly reduced in pups from infected mothers compared to their uninfected counterparts, with reductions in both effector (IL-7R^low^) and memory (IL-7R^hi^) Th2 compartments (Figure 5 C,). This correlated with a marked defect in CD21/35 expression in stromal (mesenchymal) cells (Figure 5 D) and overall reduced frequency and number of FDCs in the draining lymph node (Figure 5 E).

**Figure 5.**
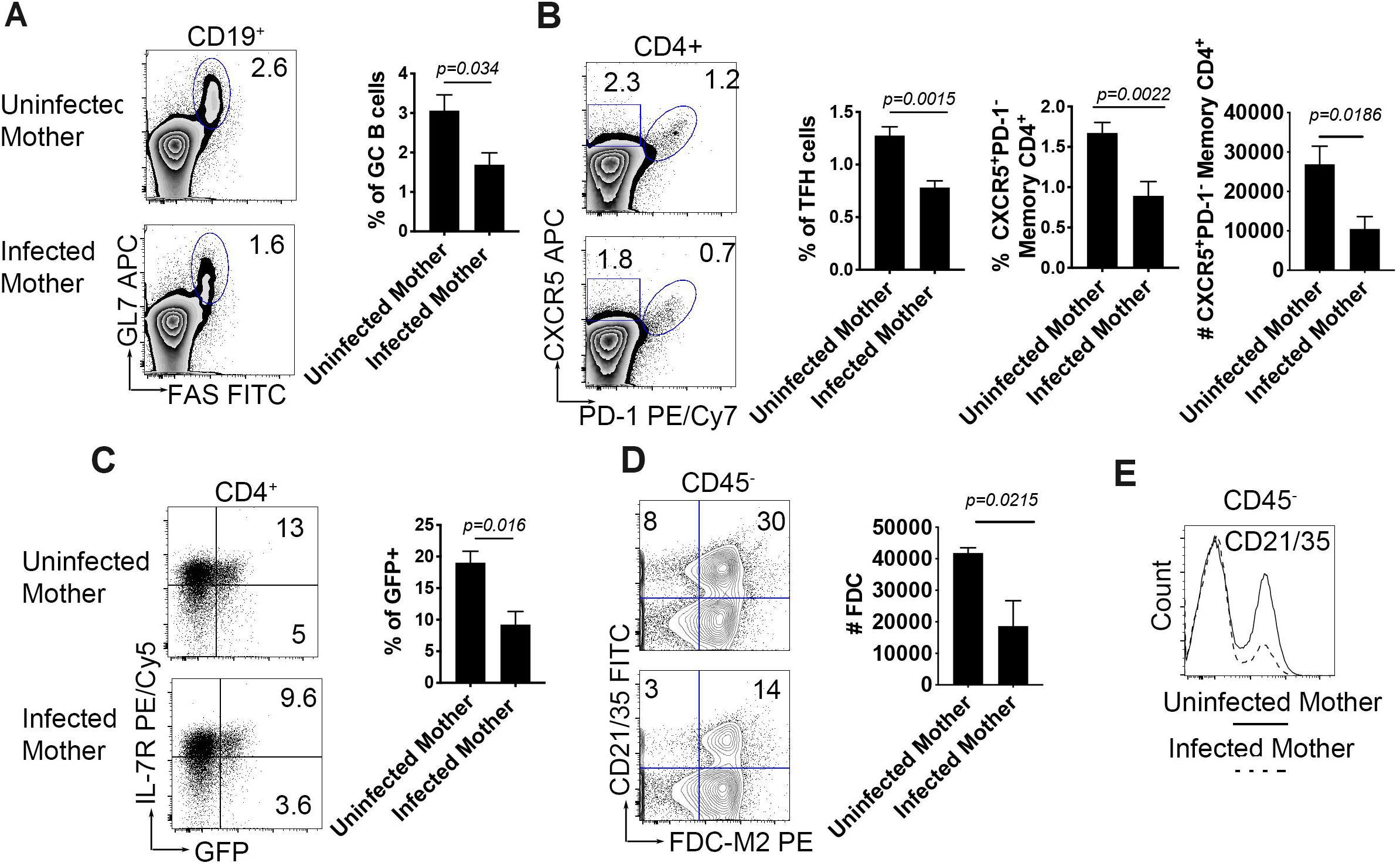
Cellular T and B responses are reduced in 4get/KN2 offspring from infected mothers 60 days post-primary immunization. 4get/KN2 28-35 days old were immunized in the hind footpad as described in the Materials and Methods. Draining popliteal lymph nodes were collected >60 days post primary immunization (A) GC B cells (CD19^+^GL-7^+^FAS^+^) frequencies (B) Tfh (PD-1^+^CXCR5^+^, gated from CD4^+^) frequencies, (C) Th2 responses (GFP^+^IL-7R ^+/-^) (D) Flow cytometry analysis of CD21/35 cell count (E) FDC (CD21/35^+^FDC-M2^+^CD45^-^) frequency. Flow data is concatenated representative of n n>4 mice per group and two independent experiments. Statistical analysis was calculated with Student’s t-test.

### Maternal schistosomiasis transcriptomically modulates the B cell lineage in an egg antigen-dependent manner

In order to more fully understand the molecular effects of maternal schistosomiasis we performed single cell RNASeq (scRNASeq) using the 10x Genomics platform. Offspring from infected, uninfected, and single-sex infected mothers (6-8 offspring per group) were immunized with tetanus/diphtheria at 28 days of age. On day 8 post-immunization live CD45^+^ immune cells were sorted from the pooled draining lymph nodes of each group. We aggregated data from 13,437 individual cells (4,879 cells-uninfected mother; 4,388 cells-infected mother; 4,170 cells-single-sex infected mother) and performed unsupervised clustering analysis based on the similarity of gene expression signatures by using the Seurat single-cell genomics R package [21]. This analysis revealed 23 distinct cell clusters representative of both lymphoid and myeloid lineages, and cluster identity was determined using CIPR [22]. For cell clusters in which the algorithm could not make a clear call such as TFH cells, we resorted to differential expression analysis between clusters to identify distinguishing markers (Supplementary Figure 1 A). Patent maternal infection (Group B) shifted the cellular proportions of the CD45^+^ pool, reducing germinal center B and TFH cells, while increasing CD8^+^ and naive CD4^+^ cells (Supplementary figure 1 B), the cellular proportions in offspring born to single-sex infected mothers is closer to that of uninfected mothers than patent infection. Upon confidently identifying cell clusters, we analyzed the differences between offspring exposed to egg antigens via patent infection and those unexposed or exposed only to adult antigens (single-sex infected, group C). Since our other experiments found a profound defect in the B cells response to immunization we focused on the follicular and germinal center clusters. Offspring born to patently infected mothers had a shared modulation of the key B cell lineage transcription factor Ebf-1, as well as multiple genes involved in cell-cycle/proliferation. The follicular B cell clusters shared a similar transcriptional profile (Supplementary Figure 2 and Figure 6 A), with follicular B cell cluster 2 having a significant reduction in Ebf-1 (adj p= 1.83e-59), a significant increase in Jun (adj p= 2.89e-18) and Junb (adj p= 3.92e-15), and significant decreases in Rbx-1 (adj p= 1.68e-13), Snrpe (adj p= 1.75e-26), Snrpg (adj p= 5.75e-83) and Tmsb10 (adj p= 3.55e-149). The germinal center B cell cluster had reduced expression of Ebf-1 (adj p= 7.05e-14), and significant decreases in Snrpe (adj p= 3.27e-20), Snrpg (adj p= 4.57e-30), Rbx1 (adj p= 8.15e-7), and Tmsb10 (adj p= 6.95e-32). Importantly, offspring born to single-sex infected mothers have no significant alterations in these genes in any B cell cluster, indicating that transcriptional alteration of cell cycle and B cell identity is dependent on egg antigen exposure, which supports our data correlating maternal titer SEA to offspring titer SEA, and offspring anti-SEA titer to offspring anti-diphtheria titer. We were unable to identify a distinct plasma cell cluster in the single-cell data, so we measured EBf-1 and cell cycle via flowcytometry. At steady-state (28-35 days of age), offspring born to patently infected mothers have reduced Ebf-1 expression in plasma cells (Figure 6 C). Ki67 is a widely used marker of active proliferation [23] so quantified Ki-67 expression in steady-state and found that a significantly lower percentage of plasma cells from offspring born to infected mother are Ki-67^+^ (Figure 6 C), suggesting reduced proliferation or cell cycle progression in these cells. We then measured Ebf-1 and Ki-67 at day 14 post immunization (the peak of the lymph node B cell response) and found that both germinal center and plasma cells in offspring born to infected mothers have reduced EBF-1 (Figure 6 D), and plasma cells have reduced Ki-67 (Figure 6 D). Indeed, when we measure the absolute number of plasma cells at steady-state and Day 14 post-immunization, we find that lymph node plasma cells in offspring from uninfected mother expand, but the offspring from infected mother fail to expand the plasma cell compartment in response to tetanus and diphtheria antigens (Figure 6 E). These data strongly support the conclusion that in utero exposure to schistosome egg antigens modulates the cell cycle and proliferative capacity of B cells, resulting in deficient plasma cell production to both microbiota and immunization.

**Figure 6.**
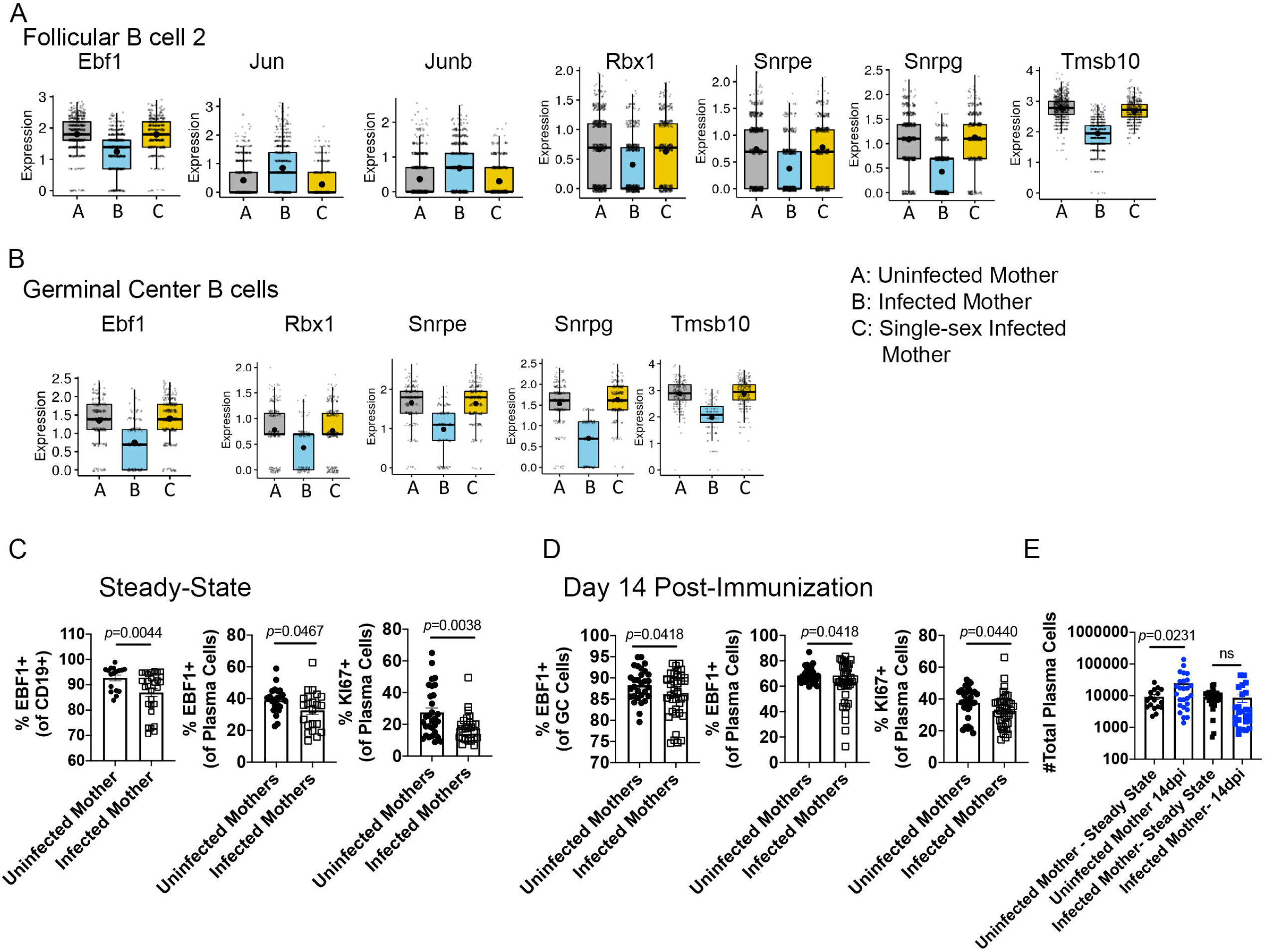
Maternal schistosomiasis induced modulation of B cell cell-cycle and proliferation is dependent on egg antigen exposure. (A,B) 4get homozygous females were infected with a low dose of *S. mansoni* cercariae to produce either single-sex infection (verified by anti-SEA ELISA and perfusion of the mother), or mixed-sex patent infection. Offspring born to mothers with either patent (egg exposure), single-sex (adult antigen exposure only), or control uninfected mothers were immunized at 28 days of age with commercial alum adjuvanted Tetanus/Diphtheria. At 8 days post-immunization CD45^+^live cells were sorted and processed for 10×Genomics Single-cell sequencing. Expression levels of genes significantly altered in follicular B cluster 2 and germinal B cell clusters. In these plots, each dot represents a single cell. Normalized expression values were used, and random noise was added to show the distribution of data points. The box plots show interquartile range and the median value (bold horizontal bar). The average expression value per sample is indicated by the dot. Wilcoxon’s test was used for statistical comparisons. (C) Flow cytometry analysis of Ebf-1 and Ki-67 at steady state in CD19+ and plasma B cells. (D) Flow cytometry analysis of Ebf-1 and Ki-67 at day 14 post tetanus/diphtheria immunization. (E) Total number of plasma cells in the popliteal lymph node at steady state and Day 14 post tetanus/diphtheria immunization. Statistical analysis for C-E was calculated with Student’s t-test with Welch’s correction.

## Discussion

In this study we report the generation of a maternal *Schistosoma mansoni* infection model in dual IL-4 reporter mice. Our results indicate that *in utero* egg antigen sensitization occurs, which is consistent to what has been published by other authors [24] and confers a state of immune-hyporesponsiveness in the offspring at steady state, this study does not differentiate between trans-placental antigen exposure and sensitization through breastfeeding; however, it has been previously suggested that exposure to parasitic antigens, either *in utero* or via breast milk, diminishes the heterologous response [24], and in some other cases nursing by infected mothers protected offspring against infection with the same helminth [25]. We observed anti-*S. mansoni* egg antigen-specific IgG1 (the dominant isotype produced against egg antigens [16]) at 35, 90 and 180 days of age in mice from infected mothers and no detectable titers in mice born to uninfected controls, suggesting either maternal antibody, antibody-secreting cell transfer to the offspring, or *in situ* antibody production by the offspring. The placental transfer of different IgG subclasses has been well documented [26-28]. In our model, we observed maternal antibodies in KN2 homozygous (IL-4 deficient) pups at 35, 60, and 90 days of age, suggesting a sustained antibody response to SEA from either maternal antibodies or cells, since our previous work has demonstrated that IL-4 deficient mice fail to mount an IgG1 response to SEA [16]. The presence of an antigen specific humoral response can potentially translate to protection, or alternatively, inhibit vaccination induced protection, as is the case for measles and other vaccinations, in humans and other mammals [29]. In humans, the persistence of maternally-derived antibodies is approximately 6 -12 months, while in other mammals it varies from 3 to 6 months, depending on the species. [30]. Nevertheless, offspring born to infected mothers have a circulating humoral response specific to *S. mansoni*, future work will determine what proportion of these cells are of maternal versus fetal origin. Previous reports have documented the presence of profibrotic biomarkers in cord blood from neonates born to mothers infected with *S. japonicum*, which was attributed to the fetal response to schistosome antigens, as the transfer of collagen markers is unlikely [31].

Interestingly, our assessment of the immune response of offspring at steady-state revealed that bulk antibody responses were impaired in offspring from infected mothers in comparison to their uninfected counterparts. This state correlated to diminished follicular dendritic cell populations in the peripheral lymph nodes. In addition to this, the area of complement receptor 1 and 2 positive FDCs in B cell follicles is also reduced at steady state and BaffR expression on B cells is reduced. Collectively, these findings are consistent with a role for FDCs in the impaired maintenance of plasma and memory B cells seen in offspring from infected mothers. FDCs are the stromal cells located in the germinal center that retain antigen for a long period and bind B cells to prevent apoptosis. These cells have previously been demonstrated to be fundamental for maintenance of plasma cells and long-lived memory B cells [32]. Recent work in our laboratory has demonstrated that IL-4 is required for FDC maintenance and positioning at steady via induction of lymphotoxin α/β [20]. In light of that, we sought to determine the cellular source of steady-state lymph node IL-4 production in this model. We found that iNKT cells are the major producers of IL-4 in control offspring during early life, and that steady-state secretion of IL-4 is almost eliminated in offspring born to infected mothers. This data suggests that the observed defects in FDC numbers and stimulatory capacity may be due to insufficient IL-4 during lymph node development and maturation. Bolstering this is the finding that there is a significant reduction in IL-4 competent (GFP^+^) CD4 T cells in the peripheral blood of these mice. This evidence suggests that prenatal helminth sensitization has a lasting effect on offspring immunity, and that offspring from mothers that were infected during pregnancy could be at a higher risk of infection with various pathogens due to a defect in homeostatic immunity as has been previously postulated [33], which will be the focus of future studies in our lab.

We further evaluated the immune response following primary immunization with a commercially available alum adjuvanted Tetanus/Diphtheria vaccine used in clinical settings. There is extensive evidence in humans of impaired response to immunization [7, 34], but little is known of the mechanism(s) controlling this defect. We observed that following primary immunization, the pups from infected mothers exhibit a defect not only in expansion of FDCs, but also in the development of T follicular helper and germinal center B cells. We hypothesize that the combined FDC and Tfh deficiency leads to the diminished response to primary immunization, as well as diminished memory T and B cell responses. Indeed, at 8-days post-immunization, there is a reduction of bulk memory B cells and IgG1^+^ B cells, which could potentially have an impact in the long-term maintenance of humoral immunity necessary for protection following vaccination. This is bolstered by our data at day 14 post-immunization where germinal center formation in a reactive lymph node was impaired in 4get/KN2 pups from infected mothers in comparison to pups to uninfected mothers. Acute antigen exposure causes lymphocyte proliferation, which is followed by a pool of quiescent long-lived IL-7R^+^ memory cells capable of potent response to challenge with the same antigen. [35]. We have previously determined that in a type 2 response, secondary TFH cells are generated in large part by recruitment of circulating Th2 memory T cells (IL-7R^+^GFP^+^) back to the lymph node, and that their interaction with memory B cells is critical to the secondary plasma cell response [16]. In our model of maternal schistosomiasis, we observed a diminished pool of both populations of IL-7R^+^GFP^+^CD4^+^ (memory Th2 cells) and IL-7R^-^GFP^+^CD4^+^ cells (Th2 effector cells) and TFH circulating precursors (CXCR5^+^PD-1^-^CD4^+^). This was accompanied by a reduction in the frequencies of Tfh cells and plasma cells, as well as a diminished secretion of IL-4 cytokine in the germinal center of the reactive lymph node (visualized in confocal tile scans). In addition to the defects observed in T cells, we also observed reduced FDC and bulk memory B cell population. Overall, this indicates that signals required for mounting a robust adaptive immune response to vaccination were dampened following prenatal exposure to *S. mansoni* antigens. In order to identify potential molecular mechanism underlying the observed B cell dysfunction we performed scRNASeq on offspring from uninfected, patently infected, and single sex infected mothers on the 10X genomics platform. Analysis of the cluster level response revealed a coordinated modulation of follicular and germinal center B cells, with profound reduction in the lineage defining transcription factor Ebf-1, and multiple genes involved in cell cycle/proliferation, such as Snpre and Snprg, which are part of the spliceosome machinery whose overexpression have been shown to drive cell proliferation in multiple models [36, 37]; Rbx1, a known modulator of DNA replication licensing proteins [38]; Tmsb10, a regulator of actin cytoskeleton with roles in B cells [39, 40]. Concomitantly, Jun and Junb, which are involved in both B cell differentiation and B cell receptor signaling [41, 42] were increased in these clusters. We validated that Ebf-1 expression is reduced in both bulk CD19^+^ and plasma cells at steady-state (28-35 days of age) and found that a significantly lower proportion of steady-state plasma cells express Ki-67, a state that persists following immunization, strongly indicating that there is indeed a defect in B cell expansion and differentiation in response to foreign antigens.

Concomitantly with defects in B cell proliferation, we find a consistent defect in follicular dendritic cell expansion and expression of C21/35. While our data suggests a role for iNKT cell IL-4 production in FDC maintenance, the data is not definitive. Future work will focus on identifying the specific homeostatic role of iNKT cell derived IL-4, and the molecular mechanism that underlie both iNKT cell production of IL-4, and diminished FDC cell function in the context of maternal infection. There is a body of literature supporting the many effects of the prenatal environment on inflammatory diseases during adolescence and adulthood. It has been hypothesized that inflammatory responses and infections during pregnancy might alter epigenetic profiles in the fetus [43]. Future studies linking the epigenetic effects of schistosomiasis infection during pregnancy and immune responses in offspring will help elucidate potential pathways involved in the immune hypo-responsiveness observed in this model.

One of the key goals of vaccination is to induce an immunological memory that is protective upon re-challenge with the same antigen [44-46]. Hence, we explored whether there was a difference between pups from infected and control mothers in the development of a sustained germinal center and maintenance of memory B and T cell pools. Pups were immunized and their immunological responses were assessed at >70 days post immunization. Surprisingly, offspring from infected mothers mounted a weaker cellular response, with significantly smaller germinal centers that correlated to a significant reduction in long lived Tfh, memory Th2 cell, and FDCs, as well as reduced complement receptor 2/1 (CD21/35) expression. Importantly, our previous work has established precursor bulk memory T cells (CXCR5^+^PD-1^-^) as critical to the secondary TFH cell response, and this cell population was significantly reduced at the memory time-points in pups from infected mothers in comparison to pups from uninfected mothers. These data strongly support the premise that *in utero* exposure to schistosome egg antigens transcriptionally reprograms the proliferative potential of the B cell lineage to foreign antigens, as well as the immune complex presenting capacity of FDCs, leading to long-lived suppression of the stromal-lymphocyte axis and the ability to mount a B cell response to immunization.

## Acknowledgements

KCF conceived of the project, KCF and DCS designed the experiments, KCF, DCS, LG, and AR performed experiments, KCF, DCS, and BR analyzed the data, KCF, and DCS wrote the manuscript. The work was supported by The University of Utah, a Scientist Development Grant from the American Heart Association to KCF (14SDG18230012), an R01 (AI135045) to KCF and an American Heart Association Award (18PRE34030086) to DCS. *B. glabrata* snails provided by the NIAID Schistosomiasis Resource Center of the Biomedical Research Institute (Rockville, MD) through NIH-NIAID Contract HHSN272201700014I for distribution through BEI Resources.

## Experimental models and subject details

### Mice strains and in vivo treatments

4get homozygous (Il4^tm1Lky^, The Jackson Laboratory), KN2 homozygous (Il4^tm1(CD2)Mmrs^, gift from Markus Morhs [15]) and 4get/KN2 were bred in-house under specific pathogen-free (SPF) conditions at Purdue University and the University of Utah. Experimental procedures involving mice were approved by the University of Utah Animal Care and Use Committee. Homozygous 4get or heterozygous 4get/KN2 females of 7-8 weeks of age were either infected with a low dose (30-35 cercariae) of *Schistosoma mansoni* or mock-infected. Infection was performed by exposing mice percutaneously to the parasite cercariae for 25 minutes. Infected and uninfected females (as controls) were bred with naive KN2 homozygous male mice at four weeks post-infection and resulting 4get/KN2 pups were weaned and genotyped (as necessary) at 28 days old. A mix of age-matched female and male pups either from control or infected females were sacrificed at 29-35 days old for steady state studies. Other experimental mice were immunized with 1/10^th^ of the human dose of Tetanus/Diphtheria commercial vaccine (Tetanus and Diphtheria Toxoids Adsorbed, MassBiologics, Boston, Massachusetts) subcutaneous (s.c.) in the rear footpad and mice were sacrificed at 8, 14, and over 60 (memory) days post-immunization.

## Method details

### Isolation of cells and flow cytometry

Cells were isolated from popliteal lymph node (pLN), hepatic lymph node (hLN), whole blood, and spleen. For follicular dendritic cells (FDC) analysis, tissues were digested as previously described ([20, 47]. Briefly, lymph nodes were digested at 37°C for 20 minutes, with occasional inversion of the contents and filtered through 100 μm filters. Single-cell suspensions were stained with surface markers and intracellular staining was performed as previously described [48]. Antibodies conjugated with the following fluorochromes were used: allophycocyanin, allophycocyanin-Cy7, Pacific blue, Brilliant Violet 510, Brilliant Violet 605, Brilliant Violet 650, Brilliant Violet 711, Brilliant Violet 786, Super Bright 600, FITC, Phycoerythrin, PE/Cy7, PE/Cy5, efluor660, PerCpCy5.5, APCAlexafluor700, and Zombie Red. The following antibodies from eBioscience, BD Biosciences, Biolegend were used: CD3 (17A2), CD4 (RM4-5), CD19 (1D3), CD138, (281-2), IgG1 (A85-1), IgD (11-26), GL7 (GL7), CD45 (30-F11), CD31 (390), podoplanin (eBIO 8.1.1), FDC-M2 (FDC-M2), CD21/35 (7G6), PD-1 (J43), CXCR5 (2G8), TCRβ (H57-597), DX5 (CD49b), CD27 (LG.3A10), IFNγ (XMG1.2), IL-7R (SB1199), FAS (15A7), HuCD2 (RPA-2.10), Ki-67 (sola15), Ebf-1 (T26-818), BaffR (7H22-E16), CD38 (90), and IL-21R/FC chimera (R&D Systems; 596-MR-100). Tetramer staining was performed using Mouse CD1d-tetramer loaded with the α-GalCer analog PBS57 and unloaded mouse CD1d-tetramer was used as staining control. Both loaded and unloaded tetramers were provided by the NIH Tetramer Core Facility.

### Immunofluorescence microscopy

Whole tissue (popliteal lymph node or hepatic lymph node) was collected and placed in Tissue-Tek optimum cutting temperature compound (OCT) (Thermo Scientific) and frozen in liquid nitrogen. Serial cryostat sections (10μm) were collected using a Leica CM 1850. Sections were then air-dried and fixed in ice-cold 75% acetone/25% ethanol for 5 mins. Sections were rehydrated in PBS for 5 to 10 minutes and blocked using biotin blocking kit (Vector Laboratories) followed by incubation with 1%(v/v) in PBS of rat and rabbit serum. Staining with appropriate antibodies was done overnight at 4°C followed by secondary staining for 1 hour at room temperature. Coverslips were mounted using ProLong anti-fade reagents (Life Technologies). Images were acquired with a Leica TCS Sp5 Laser Scanning Microscopy with an average grid size of 3×3. Images were taken with a 20x objective at a resolution of 1024×1024. Image post-processing was done using Fiji is Just ImageJ software (1.47v).

### ELISA

Schistosoma mansoni egg antigens (SEA) and tetanus and diphtheria-specific IgG1 endpoint titers were determined by enzyme-Linked immunosorbent assay (ELISA) using the mAb X56 (BD) and Immulon 4HB plates (Thermo Fisher Scientific) as previously described [20]. Briefly, plates were coated with 2μg/mL tetanus (List Labs), diphtheria (Sigma), or SEA. The following morning plates were blocked with 1% milk and incubated with serial dilutions of primary antibody, followed by incubation with anti-mouse IgG1 ads-HRP antibody (Southern Biotech) and ABTS substrate. Plates were read at 405 nm at room temperature on a BioTek plate reader.

## Single Cell RNASeq

### Sample preparation and sequencing

Single-cell suspension form 6-8 draining lymph nodes per group were pooled and stained with Dapi (Sigma) and anti-CD45. Live CD45+ cells were sorted via BD FACSAria cell sorter and washed once in PBS containing 0.04% BSA. Samples were then processed for SCseq via a 10× platform according to the manufacturer’s instructions (10× Genomics). Paired-end RNASeq (125 cycles) was performed via an Agilent HiSeq next-generation sequencer. Sequencing reads were processed by using 10× Genomics CellRanger pipeline and further analyzed with the Seurat R package. The effect of mitochondrial gene representation and the variance of unique molecular identifier (UMI) counts were regressed out from the data set prior to analysis. Gene expression signatures defining cell clusters were analyzed after aggregating 3 samples (uninfected mother, infected mother, single-sex infected mother). The raw data from scRNASeq experiments in this manuscript can be found in the NCBI’s Gene Expression Omnibus database (TBD).

#### Identification of cell clusters

Cells in our data set were clustered by using the FindClusters function of the Seurat analysis package, which identifies clusters via a shared nearest neighbor (SNN) modularity optimization– based algorithm. This function identified 23 distinct clusters spanning the lymphoid and myeloid cell lineages. The biological identities of cell clusters were annotated with the help of an immune-cell scoring algorithm ([22]and available at http://labs.path.utah.edu/oconnell/resources.htm) and by surveying known immune cell markers in the SCseq data set. The immune-scoring algorithm compares the gene signatures of the cell clusters in this study with the publicly available microarray data hosted in the Immunological Genome Project Database (ImmGen). By using differentially expressed gene signatures from Seurat, the immune-scoring algorithm performs the following steps: (a) for each ImmGen cell population, and for each gene found in ImmGen microarrays, it calculates the ratio of normalized microarray signal to the average signal value of the gene from the whole ImmGen data; (b) applies natural log transformation to the ratio, resulting in positive numbers for upregulated genes and negative numbers for downregulated genes in ImmGen data sets; (c) multiplies ImmGen log-ratio values with the log-ratio of matching genes that are differentially expressed in each cell cluster in the SCseq dataset; and (d) sums up scores from all the genes to yield an aggregate identity score for each ImmGen cell type for a given SCseq cluster. In this approach, genes that are differentially upregulated or downregulated in both ImmGen and SCseq data sets contribute to the immune identity score more heavily (a positive number is obtained when 2 log-ratio values with the same sign are multiplied). In contrast, if a gene is inversely regulated in ImmGen and SCseq clusters, the immune identity score is reduced. Through this method, the correlation between the gene expression signatures of SCseq cell clusters in our study and ImmGen data subsets assists in determining the cluster identities. In cases where this algorithm is unable to make a clear call (as T follicular helper cells and myeloid cells), we surveyed the expression of known genes in the data set and performed differential expression analyses between closely related cell clusters. This approach allowed us to further differentiate subsets. Upon naming the clusters, the Seurat R package was used to create plots for the expression of selected genes. GSEA analysis was performed by using fgsea R package [49], after ranking genes using a signal-to-noise metric [50].

### Statistical analysis

Statistical analyses were performed using either a non-parametric Mann-Whitney test, unpaired Student’s t-test with Welch’s correction, or ANOVA with Bonferroni’s multiple comparison test based on the distribution of the data. p values ≤ 0.05 were considered statistically significant. To test the strength of the association between variables, Pearson correlation coefficient was calculated. Graph generation and statistical analyses were performed using Prism (GraphPad v8.0).

## References

1. Olveda DU, Olveda RM, McManus DP, Cai P, Chau TN, Lam AK, et al. The chronic enteropathogenic disease schistosomiasis. Int J Infect Dis. 2014;28:193-203. Epub 2014/09/25. doi: 10.1016/j.ijid.2014.07.009. PubMed PMID: 25250908.

2. Steinmann P, Keiser J, Bos R, Tanner M, Utzinger J. Schistosomiasis and water resources development: systematic review, meta-analysis, and estimates of people at risk. Lancet Infect Dis. 2006;6(7):411-25. Epub 2006/06/23. doi: 10.1016/S1473-3099(06)70521-7. PubMed PMID: 16790382.

3. Colley DG, Bustinduy AL, Secor WE, King CH. Human schistosomiasis. Lancet. 2014;383(9936):2253-64. Epub 2014/04/05. doi: 10.1016/S0140-6736(13)61949-2. PubMed PMID: 24698483; PubMed Central PMCID: PMCPMC4672382.

4. McDonald EA, Pond-Tor S, Jarilla B, Sagliba MJ, Gonzal A, Amoylen AJ, et al. Schistosomiasis japonica during pregnancy is associated with elevated endotoxin levels in maternal and placental compartments. The Journal of infectious diseases. 2014;209(3):468–72. doi: 10.1093/infdis/jit446. PubMed PMID: 23964108; PubMed Central PMCID: PMC3883168.

5. Siegrist D, Siegrist-Obimpeh P. Schistosoma haematobium infection in pregnancy. Acta Trop. 1992;50(4):317-21. Epub 1992/04/01. PubMed PMID: 1356302.

6. Kurtis JD, Higashi A, Wu HW, Gundogan F, McDonald EA, Sharma S, et al. Maternal Schistosomiasis japonica is associated with maternal, placental, and fetal inflammation. Infection and immunity. 2011;79(3):1254–61. doi: 10.1128/IAI.01072-10. PubMed PMID: 21149589; PubMed Central PMCID: PMC3067505.

7. Ondigo BN, Muok EMO, Oguso JK, Njenga SM, Kanyi HM, Ndombi EM, et al. Impact of Mothers’ Schistosomiasis Status During Gestation on Children’s IgG Antibody Responses to Routine Vaccines 2 Years Later and Anti-Schistosome and Anti-Malarial Responses by Neonates in Western Kenya. Front Immunol. 2018;9:1402. Epub 2018/07/04. doi: 10.3389/fimmu.2018.01402. PubMed PMID: 29967622; PubMed Central PMCID: PMCPMC6015899.

8. Malhotra I, Mungai P, Wamachi A, Kioko J, Ouma JH, Kazura JW, et al. Helminth- and Bacillus Calmette-Guerin-induced immunity in children sensitized in utero to filariasis and schistosomiasis. J Immunol. 1999;162(11):6843-8. PubMed PMID: 10352306.

9. Elias D, Akuffo H, Pawlowski A, Haile M, Schon T, Britton S. Schistosoma mansoni infection reduces the protective efficacy of BCG vaccination against virulent Mycobacterium tuberculosis. Vaccine. 2005;23(11):1326-34. Epub 2005/01/22. doi: 10.1016/j.vaccine.2004.09.038. PubMed PMID: 15661380.

10. Attallah AM, Abbas AT, Dessouky MI, El-emshaty HM, Elsheikha HM. Susceptibility of neonate mice born to Schistosoma mansoni-infected and noninfected mothers to subsequent S. mansoni infection. Parasitology research. 2006;99(2):137–45. doi: 10.1007/s00436-006-0127-x. PubMed PMID: 16521039.

11. Santos P, Sales IR, Schirato GV, Costa VM, Albuquerque MC, Souza VM, et al. Influence of maternal schistosomiasis on the immunity of adult offspring mice. Parasitology research. 2010;107(1):95–102. doi: 10.1007/s00436-010-1839-5. PubMed PMID: 20372927.

12. Desowitz RS, Elm J, Alpers MP. Plasmodium falciparum-specific immunoglobulin G (IgG), IgM, and IgE antibodies in paired maternal-cord sera from east Sepik Province, Papua New Guinea. Infect Immun. 1993;61(3):988-93. Epub 1993/03/01. PubMed PMID: 8432619; PubMed Central PMCID: PMCPMC302830.

13. Friedman JF, Mital P, Kanzaria HK, Olds GR, Kurtis JD. Schistosomiasis and pregnancy. Trends Parasitol. 2007;23(4):159-64. Epub 2007/03/06. doi: 10.1016/j.pt.2007.02.006. PubMed PMID: 17336160.

14. Mohrs M, Shinkai K, Mohrs K, Locksley RM. Analysis of type 2 immunity in vivo with a bicistronic IL-4 reporter. Immunity. 2001;15(2):303-11. Epub 2001/08/25. PubMed PMID: 11520464.

15. Mohrs K, Wakil AE, Killeen N, Locksley RM, Mohrs M. A two-step process for cytokine production revealed by IL-4 dual-reporter mice. Immunity. 2005;23(4):419-29. Epub 2005/10/18. doi: 10.1016/j.immuni.2005.09.006. PubMed PMID: 16226507; PubMed Central PMCID: PMC2826320.

16. Fairfax KC, Everts B, Amiel E, Smith AM, Schramm G, Haas H, et al. IL-4-secreting secondary T follicular helper (Tfh) cells arise from memory T cells, not persisting Tfh cells, through a B cell-dependent mechanism. J Immunol. 2015;194(7):2999-3010. Epub 2015/02/26. doi: 10.4049/jimmunol.1401225. PubMed PMID: 25712216; PubMed Central PMCID: PMCPMC4495582.

17. Tew JG, Wu J, Fakher M, Szakal AK, Qin D. Follicular dendritic cells: beyond the necessity of T-cell help. Trends Immunol. 2001;22(7):361-7. Epub 2001/06/29. PubMed PMID: 11429319.

18. Suzuki K, Grigorova I, Phan TG, Kelly LM, Cyster JG. Visualizing B cell capture of cognate antigen from follicular dendritic cells. The Journal of experimental medicine. 2009;206(7):1485–93. doi: 10.1084/jem.20090209. PubMed PMID: 19506051; PubMed Central PMCID: PMCPMC2715076.

19. Wu Y, Sukumar S, El Shikh ME, Best AM, Szakal AK, Tew JG. Immune complex-bearing follicular dendritic cells deliver a late antigenic signal that promotes somatic hypermutation. J Immunol. 2008;180(1):281-90. Epub 2007/12/22. PubMed PMID: 18097029.

20. Cortes-Selva D, Ready A, Gibbs L, Rajwa B, Fairfax KC. IL-4 promotes stromal cell expansion and is critical for development of a type-2, but not a type 1 immune response. European journal of immunology. 2019;49(3):428-42. Epub 2018/12/24. doi: 10.1002/eji.201847789. PubMed PMID: 30575951.

21. Butler A, Hoffman P, Smibert P, Papalexi E, Satija R. Integrating single-cell transcriptomic data across different conditions, technologies, and species. Nature biotechnology. 2018;36(5):411-20. Epub 2018/04/03. doi: 10.1038/nbt.4096. PubMed PMID: 29608179; PubMed Central PMCID: PMCPMC6700744.

22. Ekiz HA, Conley CJ, Stephens WZ, O’Connell RM. CIPR: a web-based R/shiny app and R package to annotate cell clusters in single cell RNA sequencing experiments. BMC Bioinformatics. 2020;21(1):191. Epub 2020/05/18. doi: 10.1186/s12859-020-3538-2. PubMed PMID: 32414321; PubMed Central PMCID: PMCPMC7227235.

23. Li X, Miao H, Henn A, Topham DJ, Wu H, Zand MS, et al. Ki-67 expression reveals strong, transient influenza specific CD4 T cell responses after adult vaccination. Vaccine. 2012;30(31):4581-4. Epub 2012/05/05. doi: 10.1016/j.vaccine.2012.04.059. PubMed PMID: 22554464; PubMed Central PMCID: PMCPMC3858959.

24. Santos P, Lorena VM, Fernandes Ede S, Sales IR, Nascimento WR, Gomes Yde M, et al. Gestation and breastfeeding in schistosomotic mothers differently modulate the immune response of adult offspring to postnatal Schistosoma mansoni infection. Memorias do Instituto Oswaldo Cruz. 2016;111(2):83–92. doi: 10.1590/0074-02760150293. PubMed PMID: 26872339; PubMed Central PMCID: PMCPMC4750447.

25. Matthew G. Darby AC, Dunja Mrjden, Marion Rolot, Katherine Smith, Claire Mackowiak, Delphine Sedda, Donald Nyangahu, Heather Jaspan, Kai-Michael Toellner, Ari Waisman, Valerie Quesniaux, Bernhard Ryffel, Adam F. Cunningham, Benjamin G. Dewals, Frank Brombacher, William G. C. Horsnell. Pre-conception maternal helminth infection transfers via nursing long-lasting cellular immunity against helminths to offspring. Science Advances. 2019;5(5). doi: 10.1126/sciadv.aav3058

26. Lostal Gracia MI, Larrad Mur L, Perez Gonzalez JM. [IgG subclasses: placental transfer in the full-term neonate and their evolution during the first 3 months of life]. An Esp Pediatr. 1993;38(6):503-8. Epub 1993/06/01. PubMed PMID: 8368678.

27. Okoko BJ, Wesumperuma LH, Ota MO, Pinder M, Banya W, Gomez SF, et al. The influence of placental malaria infection and maternal hypergammaglobulinemia on transplacental transfer of antibodies and IgG subclasses in a rural West African population. J Infect Dis. 2001;184(5):627-32. Epub 2001/08/09. doi: 10.1086/322808. PubMed PMID: 11494168.

28. Pitcher-Wilmott RW, Hindocha P, Wood CB. The placental transfer of IgG subclasses in human pregnancy. Clin Exp Immunol. 1980;41(2):303-8. Epub 1980/08/01. PubMed PMID: 7438556; PubMed Central PMCID: PMCPMC1537014.

29. Niewiesk S. Maternal antibodies: clinical significance, mechanism of interference with immune responses, and possible vaccination strategies. Front Immunol. 2014;5:446. Epub 2014/10/04. doi: 10.3389/fimmu.2014.00446. PubMed PMID: 25278941; PubMed Central PMCID: PMCPMC4165321.

30. Tsutsui T, Yamamoto T, Hayama Y, Akiba Y, Nishiguchi A, Kobayashi S, et al. Duration of maternally derived antibodies against Akabane virus in calves: survival analysis. J Vet Med Sci. 2009;71(7):913-8. Epub 2009/08/05. PubMed PMID: 19652478.

31. McDonald EA, Cheng L, Jarilla B, Sagliba MJ, Gonzal A, Amoylen AJ, et al. Maternal infection with Schistosoma japonicum induces a profibrotic response in neonates. Infect Immun. 2014;82(1):350-5. Epub 2013/10/30. doi: 10.1128/IAI.01060-13. PubMed PMID: 24166958; PubMed Central PMCID: PMCPMC3911825.

32. Al-Alwan M, Du Q, Hou S, Nashed B, Fan Y, Yang X, et al. Follicular dendritic cell secreted protein (FDC-SP) regulates germinal center and antibody responses. J Immunol. 2007;178(12):7859-67. Epub 2007/06/06. PubMed PMID: 17548624.

33. Lacorcia M, Prazeres da Costa CU. Maternal Schistosomiasis: Immunomodulatory Effects With Lasting Impact on Allergy and Vaccine Responses. Front Immunol. 2018;9:2960. Epub 2019/01/09. doi: 10.3389/fimmu.2018.02960. PubMed PMID: 30619318; PubMed Central PMCID: PMCPMC6305477.

34. Labeaud AD, Malhotra I, King MJ, King CL, King CH. Do antenatal parasite infections devalue childhood vaccination? PLoS neglected tropical diseases. 2009;3(5):e442. doi: 10.1371/journal.pntd.0000442. PubMed PMID: 19478847; PubMed Central PMCID: PMC2682196.

35. Seddon B, Tomlinson P, Zamoyska R. Interleukin 7 and T cell receptor signals regulate homeostasis of CD4 memory cells. Nature immunology. 2003;4(7):680-6. Epub 2003/06/17. doi: 10.1038/ni946. PubMed PMID: 12808452.

36. Anchi T, Tamura K, Furihata M, Satake H, Sakoda H, Kawada C, et al. SNRPE is involved in cell proliferation and progression of high-grade prostate cancer through the regulation of androgen receptor expression. Oncol Lett. 2012;3(2):264-8. Epub 2012/06/29. doi: 10.3892/ol.2011.505. PubMed PMID: 22740892; PubMed Central PMCID: PMCPMC3362443.

37. Lan Y, Lou J, Hu J, Yu Z, Lyu W, Zhang B. Downregulation of SNRPG induces cell cycle arrest and sensitizes human glioblastoma cells to temozolomide by targeting Myc through a p53-dependent signaling pathway. Cancer Biol Med. 2020;17(1):112-31. Epub 2020/04/17. doi: 10.20892/j.issn.2095-3941.2019.0164. PubMed PMID: 32296580; PubMed Central PMCID: PMCPMC7142844.

38. Jia L, Bickel JS, Wu J, Morgan MA, Li H, Yang J, et al. RBX1 (RING box protein 1) E3 ubiquitin ligase is required for genomic integrity by modulating DNA replication licensing proteins. The Journal of biological chemistry. 2011;286(5):3379-86. Epub 2010/12/01. doi: 10.1074/jbc.M110.188425. PubMed PMID: 21115485; PubMed Central PMCID: PMCPMC3030344.

39. Tolar P. Cytoskeletal control of B cell responses to antigens. Nature reviews Immunology. 2017;17(10):621-34. Epub 2017/07/12. doi: 10.1038/nri.2017.67. PubMed PMID: 28690317.

40. Zhang X, Ren D, Guo L, Wang L, Wu S, Lin C, et al. Thymosin beta 10 is a key regulator of tumorigenesis and metastasis and a novel serum marker in breast cancer. Breast Cancer Res. 2017;19(1):15. Epub 2017/02/10. doi: 10.1186/s13058-016-0785-2. PubMed PMID: 28179017; PubMed Central PMCID: PMCPMC5299657.

41. Ohkubo Y, Arima M, Arguni E, Okada S, Yamashita K, Asari S, et al. A role for c-fos/activator protein 1 in B lymphocyte terminal differentiation. J Immunol. 2005;174(12):7703-10. Epub 2005/06/10. doi: 10.4049/jimmunol.174.12.7703. PubMed PMID: 15944271.

42. Yin Q, Wang X, McBride J, Fewell C, Flemington E. B-cell receptor activation induces BIC/miR-155 expression through a conserved AP-1 element. The Journal of biological chemistry. 2008;283(5):2654-62. Epub 2007/12/01. doi: 10.1074/jbc.M708218200. PubMed PMID: 18048365; PubMed Central PMCID: PMCPMC2810639.

43. Claycombe KJ, Brissette CA, Ghribi O. Epigenetics of inflammation, maternal infection, and nutrition. J Nutr. 2015;145(5):1109S-15S. Epub 2015/04/03. doi: 10.3945/jn.114.194639. PubMed PMID: 25833887; PubMed Central PMCID: PMCPMC4410493.

44. Bevan MJ. Understand memory, design better vaccines. Nat Immunol. 2011;12(6):463-5. Epub 2011/05/19. doi: 10.1038/ni.2041. PubMed PMID: 21587308; PubMed Central PMCID: PMCPMC3303227.

45. Sallusto F, Lanzavecchia A, Araki K, Ahmed R. From vaccines to memory and back. Immunity. 2010;33(4):451-63. Epub 2010/10/30. doi: 10.1016/j.immuni.2010.10.008. PubMed PMID: 21029957; PubMed Central PMCID: PMCPMC3760154.

46. Castellino F, Galli G, Del Giudice G, Rappuoli R. Generating memory with vaccination. Eur J Immunol. 2009;39(8):2100-5. Epub 2009/07/29. doi: 10.1002/eji.200939550. PubMed PMID: 19637203.

47. Hara T, Shitara S, Imai K, Miyachi H, Kitano S, Yao H, et al. Identification of IL-7-producing cells in primary and secondary lymphoid organs using IL-7-GFP knock-in mice. J Immunol. 2012;189(4):1577-84. Epub 2012/07/13. doi: 10.4049/jimmunol.1200586. PubMed PMID: 22786774.

48. Glatman Zaretsky A, Taylor JJ, King IL, Marshall FA, Mohrs M, Pearce EJ. T follicular helper cells differentiate from Th2 cells in response to helminth antigens. The Journal of experimental medicine. 2009;206(5):991-9. Epub 2009/04/22. doi: 10.1084/jem.20090303. PubMed PMID: 19380637; PubMed Central PMCID: PMC2715032.

49. Korotkevich G, Sukhov V, Sergushichev A. Fast gene set enrichment analysis. bioRxiv. 2019:060012. doi: 10.1101/060012.

50. Subramanian A, Tamayo P, Mootha VK, Mukherjee S, Ebert BL, Gillette MA, et al. Gene set enrichment analysis: a knowledge-based approach for interpreting genome-wide expression profiles. Proceedings of the National Academy of Sciences of the United States of America. 2005;102(43):15545-50. Epub 2005/10/04. doi: 10.1073/pnas.0506580102. PubMed PMID: 16199517; PubMed Central PMCID: PMCPMC1239896.

