## Supplementary material for "Maternal *schistosomiasis* impairs offspring IL-4 production and B cell expansion": Supplmental figures

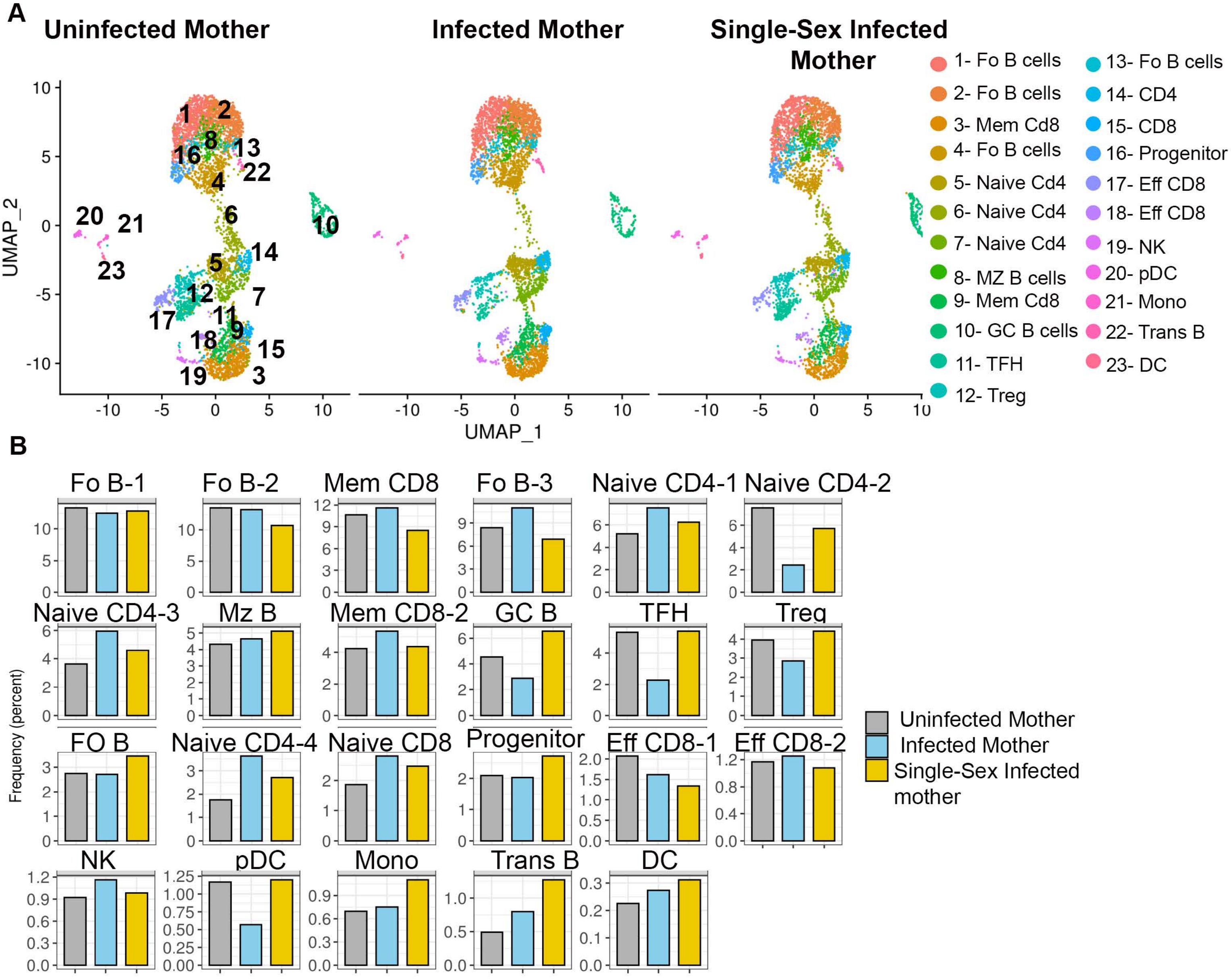


**Supplementary Figure 1**. A) scRNAseq UMAP plots showing the immune landscape in showing the immune landscape ain the draining lymph node in offspring born to uninfected , infected, and single-sex infected mothers at day 8 post tetanus/diphtheria immunization. B) Frequency of cell clusters with the indicated identity in offspring born to uninfected,infected, and single-sex infected mothers.

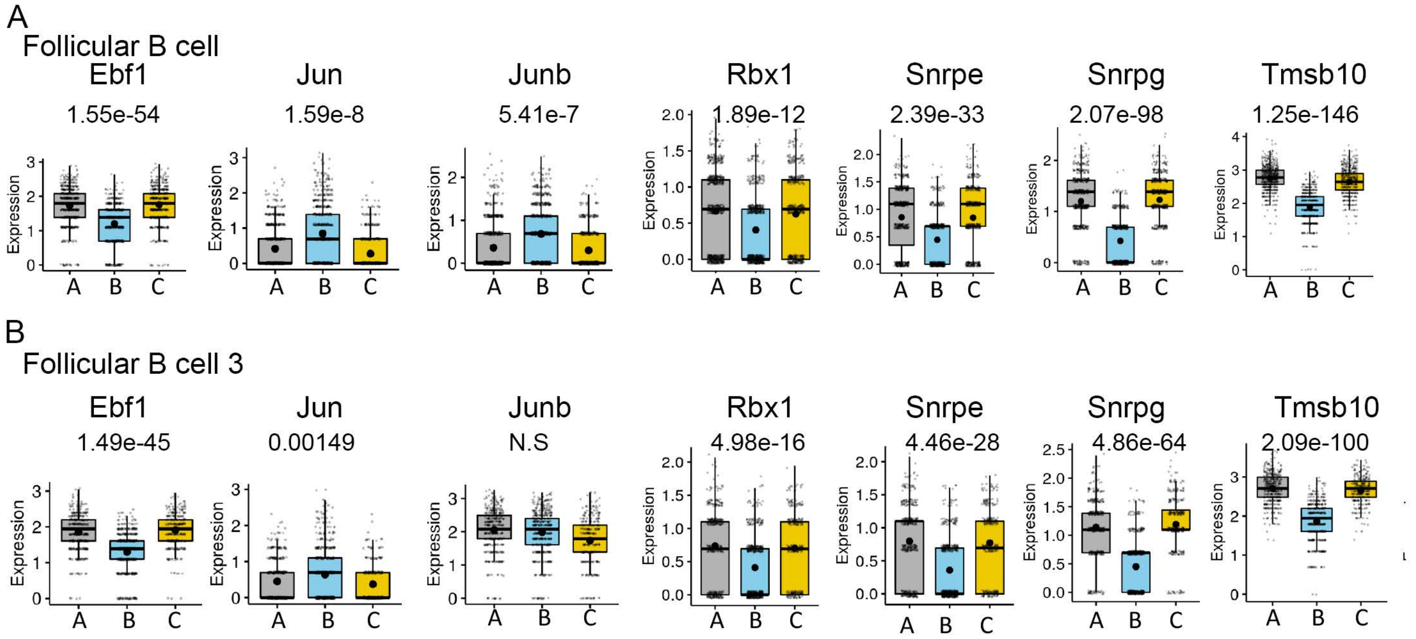


**Supplementary Figure 2**. Expression levels of Cell cycle and B cell genes genes within the Follicular B cells are shown. In these plots, each dot represents a single cell. Normalized expression values were used, and random noise was added to show the distribution of data points. The box plots show interquartile range and the median value (bold horizontal bar). Average expression value per sample is indicated by the center black points. Wilcoxon's test was used for statistical comparisons. The listed values are adjusted *p*-values.
